# Triterpenoid CDDO-EA Protects from Hyperglycemia, Hyperinsulinemia, and Obesity by Decreasing Energy Intake

**DOI:** 10.1101/2025.02.12.636924

**Authors:** Austin E. Cantu, Cordelia Rasa, Shizue Mito, Denae Cantu, Juan Carlos Lopez-Alvarenga, Leslie Rivera-Lopez, Israel Rios, Ashley Abrego-Gonzalez, Sara M. Reyna

## Abstract

Obesity is a significant factor in the development of type 2 diabetes (T2D). Treatment of obesity is pivotal in the prevention and management of T2D, and the development of new pharmacological therapies are studied for improving insulin resistance and glucose intolerance. Oleanolic acid derived triterpenoids, 2-cyano-3,12-dioxooleana-1,9-dien-28-oic acids (CDDOs), are studied to elucidate the mechanisms by which they protect against obesity. However, there remain fundamental gaps in knowledge regarding the physiological and molecular mechanisms by which CDDOs protect against obesity. Our recently published studies showed that CDDO-ethyl amide (CDDO-EA) prevents skeletal muscle inflammation by inhibiting activation of nuclear factor kappa B (NF-_κ_B) signaling. Moreover, CDDO-EA induced translocation of the glucose transporter, GLUT4, in skeletal muscle cells. We hypothesized that CDDO-EA protects from obesity-induced hyperglycemia in mice fed a high fat diet (HFD). Our results show that CDDO-EA protects from HFD-induced obesity but has no effect on body weight in mice fed a low-fat diet (LFD). Our data show that CDDO-EA inhibition of weight gain is associated with reduced caloric intake and glucose and insulin levels in mice fed a HFD. This highlights the potential of CDDO-EA as a therapeutic agent for obesity treatment and the protection against the development of T2D.

**IMPACT STATEMENT:** The significance of our studies is that they define factors affected by CDDOs to maintain energy homeostasis and improve glucose metabolism to protect against obesity-induced insulin resistance. Our work examines metabolic factors affected by CDDO-EA by demonstrating that CDDO-EA protects from weight gain, glucose intolerance, and insulin resistance. Our research adds new knowledge on the anti-obesity and anti-diabetic properties of CDDO-EA by showing that incorporation of CDDO-EA in a high fat diet prevents obesity and an increase in glucose and insulin levels in an animal model of obesity and insulin resistance. Importantly, CDDO-EA incorporated in a low-fat diet does not affect body weight and caloric intake. Given that obesity is the major risk factor in the progression to T2D, our investigation on characterizing the metabolic effects of CDDO-EA on obesity advances and facilitates the use of CDDOs as candidates for the prevention and treatment of T2D.

## INTRODUCTION

Obesity is a significant risk factor associated with the development and progression to T2D in which hyperglycemia is the primary manifestation. Furthermore, obesity is the fastest-growing pandemic in the world, and it is estimated that 738 million individuals will be diagnosed with T2D by 2045^1^. Obese individuals develop insulin resistance which is characterized by impaired insulin sensitivity by tissues^2^, leading to hyperglycemia and hyperinsulinemia which can lead to serious complications such as neuropathies, cancer, and cardiovascular disorders. Therefore, the treatment and prevention of obesity, insulin resistance, and T2D is important to prevent these serious complications.

The oleanolic acid derived triterpenoids exhibit anti-inflammatory, anti-tumorigenic, and anti-diabetic properties^3^. In particular, the CDDO derivatives, CDDO-methyl ester (CDDO-Me) and CDDO-imidazole (CDDO-Im), have been studied for their anti-diabetic properties^4-7^. Recently, we demonstrated that CDDO-EA protects skeletal muscle cells from lipopolysaccharide-induced inflammation by inhibiting NF-_κ_B activation^8^. Our work also showed that CDDO-EA induced translocation of the glucose transporter, GLUT4, in skeletal muscle cells. Our findings show that CDDO-EA protects from weight gain by preventing energy intake. Also, we show that the prevention of obesity by CDDO-EA is associated with a reduction in hyperglycemia and hyperinsulinemia. Our findings suggest that CDDO-EA holds potential as a future therapeutic agent against obesity to protect from the development of insulin resistance and T2D.

## MATERIALS AND METHODS

### Animals and Diets

All procedures were approved by the University of Texas Rio Grande Valley Institutional Animal Care and Use Committee (Protocol # 17-05). Male C57BL/6J mice (6 – 8 weeks old, strain # 000664) were purchased from the Jackson Laboratory. Mice were fed irradiated rodent diets *at libitum* as follows: low-fat diet (LFD) (10% of total calories from fat, #TD.08806, Envigo) or a high-fat diet (HFD) (60% of total calories from fat, #TD.06414, Envigo) with or without CDDO-EA (diet containing 0.04% CDDO-EA and prepared by Envigo from researcher supplied CDDO-EA) for six weeks. Mice were individually housed under controlled temperature (23°C) and lighting (10 h ligh:14 h dark) with free access to water and food. Both mice and any remaining rodent diet in each cage were weighed once a week.

### Synthesis of CDDO-EA

For our animal study, we synthesized CDDO-EA from oleanolic acid (Figure 1). First, CDDO-Me was prepared from oleanolic acid by the following the reported procedures^9^. Then, CDDO-Me was hydrolyzed in the presence of lithium iodide in dimethylformamide to produce CDDO^10^. Next, CDDO was treated with oxalyl chloride to form acid chloride, which was immediately mixed with ethylamine hydrochloride to complete the synthesis of CDDO-EA. The reaction mixture was purified by column chromatography. The structure of CDDO-EA was confirmed with 1H NMR and HRMS, and the purity was verified by HPLC (Figure S1, S2, and S3).

**Figure 1:**
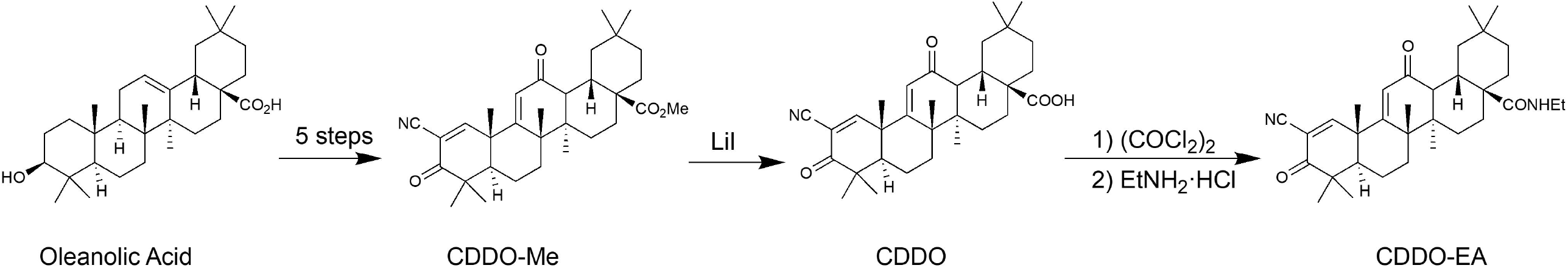
Synthesis of CDDO-EA. CDDO-Me was prepared from oleanolic acid. Next, CDDO-Me was hydrolyzed in the presence of lithium iodide in dimethylformamide to give CDDO. CDDO was treated with oxalyl chloride to form acid chloride, which was mixed with ethylamine hydrochloride to obtain CDDO-EA.

### RAW264.7 Cells

Mouse RAW 264.7 macrophage cells were cultured as described previously.^11^ The cells were sub-cultured by using 5ml of ice-cold 5mM EDTA in PBS at 4°C for 20 minutes, with the flask being tapped every 5 minutes to detach cells. The cell pellet was collected by centrifugation at 800 rpm for 5 minutes at room temperature.

### MCP-1 Detection

RAW 264.7 cells were grown on 24-well dishes at 1 × 10^6^ cells/ml, using 0.5 ml/well. Cells were pre-treated with 500 nM CDDO-EA for 1 hour. After 1 hour, cells were treated with 100 ng/ml of LPS (*Escherichia coli* O111:B4, Sigma Aldrich) for 6 hours, and supernatants were collected and stored at -80 °C until analysis. MCP-1 from the culture media of cells was measured and quantified using a Mouse MCP-1 ELISA Set (BD Bioscience, catalog # 555260) according to the manufacturer’s instructions.

### Oral Glucose Tolerance Tests and Glucose Measurements

Oral glucose tolerance tests (OGTT) were performed before the start and at week 6 of the experimental feeding. The mice were fasted for 5 hours. During OGTT, mice were awake and unrestrained. A volume of 20% dextrose solution (2 g dextrose / kg) was given to each mouse via oral gavage. Blood samples were collected via tail snips, and blood glucose levels were measured by using a glucometer. Blood glucose levels were measured at -5 minutes before glucose gavage and at 5, 10, 15, 20, 30, 45, 60, 90, and 120 minutes after glucose gavage.

Blood samples were collected via tail snips for insulin measurements at -5 minutes before glucose gavage and at 10, 30, 60, and 120 minutes after glucose gavage. Blood samples for insulin measurement were collected in blood collection tubes coated with heparin and then centrifuged for 1 minute at 13,000 rpm to collect plasma. Plasma samples were then transferred to a clean tube and stored at -80°C until ready for use. Blood glucose levels measured at -5 minutes for OGTT were used for 0- and 6-week time points. Blood samples were also collected via tail snips at 2 and 4 weeks into the experimental feeding period for blood glucose measurements using a glucometer.

### Insulin ELISA

Blood samples for measurement of insulin were collected during the OGTTs (before and at 6-week experimental feeding) and at 2 and 4 weeks into the experimental feeding. Insulin was analyzed with a Mercodia mouse insulin ELISA (catalog # 10-1247-01). ELISA was performed according to the manufacturer’s instructions.

### Statistical Analysis

All values were presented as mean ± SEM. Data were statistically analyzed by One-Way ANOVA or Two-Way ANOVA with repeated measures. *P*-values equal to or less than 0.05 were considered significant. * *P* < 0.05; \*\**P* < 0.01; *** *P* < 0.001; **** *P* < 0.0001.

## RESULTS

### Synthesized CDDO-EA Inhibits LPS-Induced MCP-1 Production in Macrophages

HPLC showed high purity (>99%) of the CDDO-EA synthesized in our lab (Figure S1, S2, and S3). Our synthesized CDDO-EA was tested for its biological property of anti-inflammatory activity. As with the original CDDO-EA which we used in our previous study and was a kind gift from Dr. Thomas Slaga from The University of Texas Health Science Center at San Antonio^8^, we tested two different batches of CDDO-EA synthesized by us for their biological activity to block MCP-1 production in macrophages. As shown in Figure 2, both batches of CDDO-EA blocked LPS-induced production of MCP-1 in RAW264.7 macrophages. This suppression of MCP-1 production shows that our synthesized CDDO-EA has anti-inflammatory properties like the original CDDO-EA^8^.

**Figure 2:**
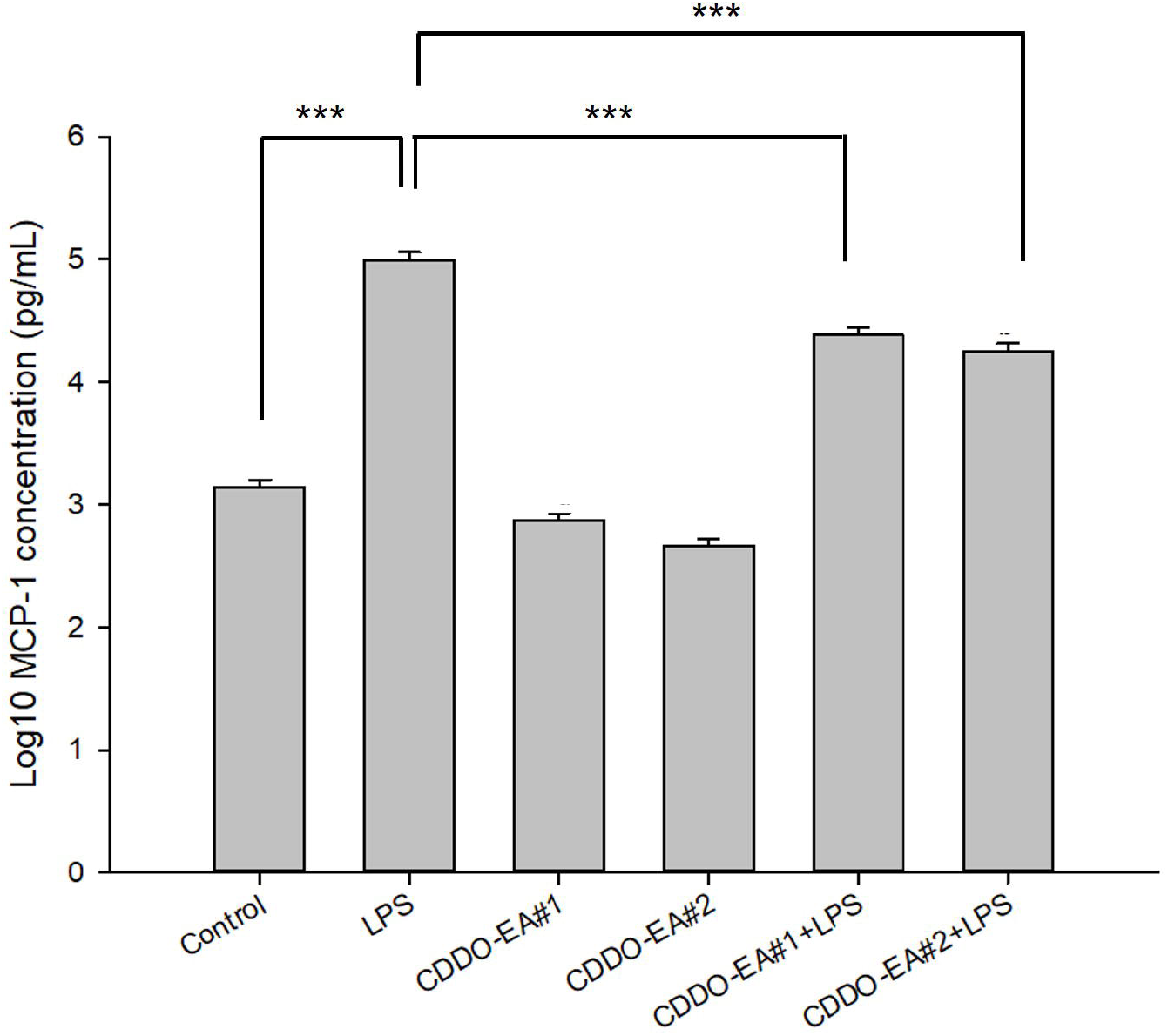
Synthesized CDDO-EA inhibits MCP-1 production in macrophages. RAW264.7 mouse macrophages were pre-treated with 500 nM CDDO-EA for 1 hour then exposed to 100 ng/ml LPS for 6 hours. MCP-1 levels were measured and quantified by ELISA. Two independent experiments were run in duplicate or triplicate. Data represented as mean ±SEM. \*\*\**p*< 0.001 (One-Way ANOVA).

### CDDO-EA Protects from High Fat Diet-Induced Obesity

As shown in Figure 3A, our findings show that mice fed a HFD weighed significantly more than the LFD fed mice by week two, and this was consistent throughout the six-week study. The incorporation of CDDO-EA in the HFD prevented excess weight gain compared to HFD alone. We then measured energy intake and found that CDDO-EA decreased energy intake in mice fed a HFD (Figure 3B). Importantly, CDDO-EA did not affect body weight and caloric intake in mice fed a LFD (Figure 3A and 3B).

**Figure 3:**
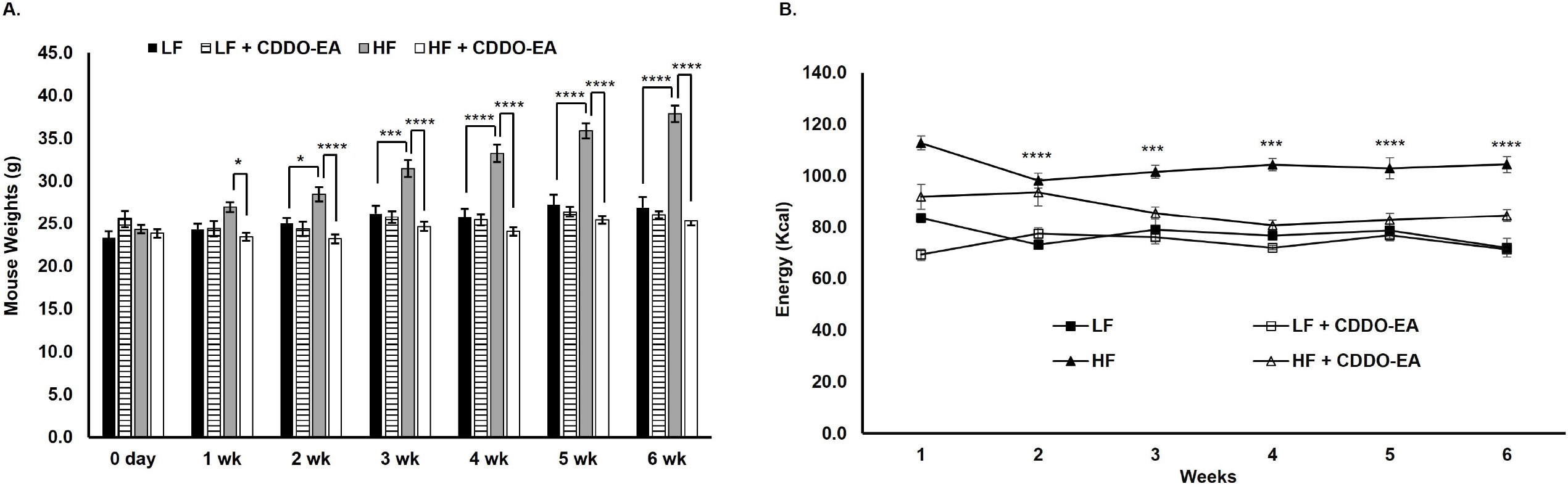
CDDO-EA prevents weight gain by inhibiting energy intake. C57Bl6/J mice were fed a LFD (n=5) or HFD (n=7) without incorporation of CDDO-EA or LFD (n=7) or HFD (n=7) with incorporation of CDDO-EA (diet containing 0.04% CDDO-EA) for 6 weeks. Mice and remaining rodent diet in each cage were weighed once a week. Data are represented as mean ± SEM; Two-way Repeated Measures ANOVA. **A**. \**p*<0.05, \*\*\**p*<0.001, \*\*\*\**p*<0.0001. **B**. \*\*\**p*<0.001 vs. HFD + CDDO-EA, \*\*\*\**p*<0.0001 vs. HFD + CDDO-EA.

### CDDO-EA Prevents Hyperglycemia and Hyperinsulinemia

We further measured glucose and insulin levels to determine if prevention of obesity by CDDO-EA coincided with a reduction in hyperglycemia and hyperinsulinemia. Blood glucose levels were significantly increased in mice fed HFD at 2 weeks and remained significantly increased throughout the rest of the period during which mice were fed the drug, as shown in Figure 4A. In contrast, blood glucose levels in mice fed a HFD with CDDO-EA did not increase throughout the six-week study. In addition, Figure 4B shows that plasma insulin levels were significantly higher at 2 weeks and remained significantly higher in mice fed only a HFD compared to mice fed a HFD with CDDO-EA. As seen in Figures 4C and 4D, OGTT showed that CDDO-EA prevented increased blood glucose and plasma insulin concentrations in mice fed a HFD. In Figure 4C, the gray shaded area corresponds to the blood glucose levels before experimental feeding. The glucose levels of both the LFD and LFD + CDDO-EA groups (left panel Figure 4C) overlap with the glucose levels before experimental feeding. Further, the HFD group’s glucose levels (right panel Figure 4C) are significantly higher after the 6-week HFD feeding. Interestingly, the HFD + CDDO-EA group’s glucose levels overlap with the HFD up to the 45-minute timepoint and then significantly decreases to glucose levels before experimental feeding. In Figure 4D, the gray shaded area corresponds to the insulin levels before experimental feeding. The insulin levels of LFD and LFD+CDDO-EA after the 6-week experimental feeding overlap with the insulin levels before experimental feeding (left panel Figure 4D). In addition, the insulin levels of the HFD group (right panel Figure 4D) are significantly higher after the 6-week HFD feeding. The HFD+CDDO-EA group insulin levels did not increase significantly and overlap with the insulin levels before experimental feeding. Our findings show that CDDO-EA did not affect glucose and insulin levels of mice fed LFD with CDDO-EA, and a feeding of HFD alone induces glucose intolerance and hyperinsulinemia. However, CDDO-EA incorporation in the HFD inhibited increases in glucose and insulin levels.

**Figure 4:**
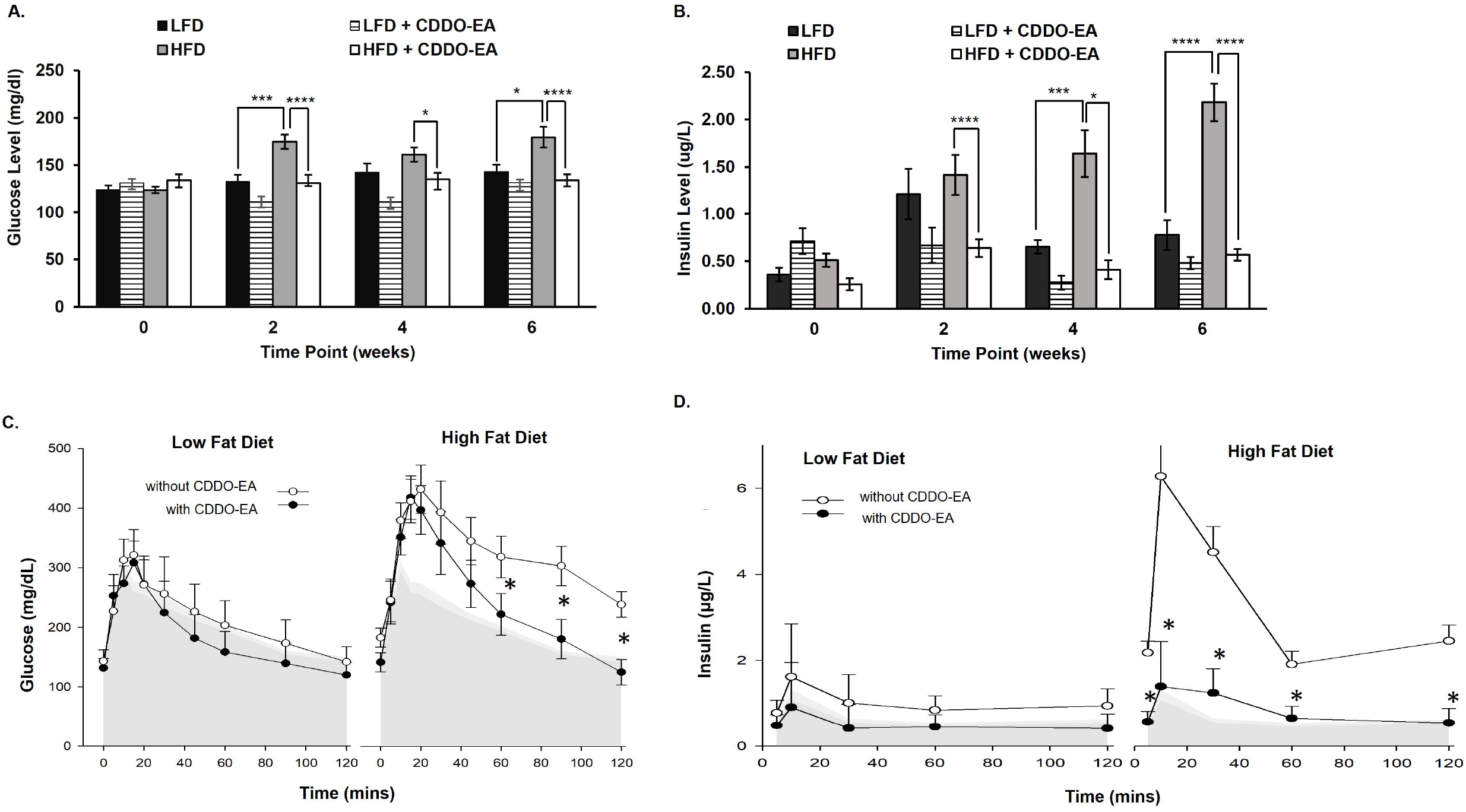
CDDO-EA prevents glucose intolerance and hyperinsulinemia. C57Bl6/J mice were fed a LFD or HFD with or without incorporation of CDDO-EA (diet containing 0.04% CDDO-EA) for 6 weeks. Blood samples were also collected via tail snips for blood glucose measurements using a glucometer and for plasma insulin measurements using an ELISA. LFD, n=5; LFD + CDDO-EA, n=7; HFD, n=7; HFD + CDDO-EA, n=7. **A and B**. Data are represented as mean ± SEM; Two-way Repeated Measures ANOVA. \**p*<0.05, \*\*\**p*<0.001, \*\*\*\**p*<0.0001. **C and D**. Blood glucose (A) and plasma insulin (B) measurements obtained from OGTTs. The left panel of each graph corresponds to LFD and the right panel of each graph corresponds to HFD. ANOVA: Differences by fat diet at basal p=0.89, final p<0.001. Differences by CDDO-EA at basal p=0.64, final p=0.003. The gray shaded area represents the mean values of glucose (A) or insulin (B) concentration (basal levels) before experimental feeding. The error bars correspond to 95% confidence intervals.

## DISCUSSION

Obesity is a high-risk factor for the development of insulin resistance, and one of the main causes of insulin resistance is impaired glucose transport in tissues, like in the skeletal muscle^2^. Skeletal muscle plays a crucial role in whole body glucose metabolism, and GLUT4 is the primary glucose transporter in skeletal muscle. Although many studies have examined anti-diabetic properties of CDDOs to protect from obesity and T2D, more studies are needed to examine their role in protecting from obesity-induced insulin resistance. This led us to study whether CDDO-EA induces GLUT4 translocation, and we previously showed that CDDO-EA indeed induced the translocation of GLUT4 in skeletal muscle cells^8^.

Thus, we wanted to investigate whether CDDO-EA protects from obesity-induced insulin resistance in an animal model. Results from our study demonstrate that CDDO-EA protects from obesity-induced insulin resistance by inhibiting weight gain due to reduced food intake. This is accompanied by improved insulin sensitivity and glucose metabolism.

CDDO-methyl ester (CDDO-Me) and CDDO-imidazole (CDDO-Im) have been mainly studied for their anti-obesity and anti-diabetic properties, and our findings add another CDDO derivative which has similar properties with some important differences. Shin, et. al. conducted a four-day indirect calorimetry study in C57BL/6J female mice after 82 days of HFD (60% calories from fat) feeding and an oral gavage of CDDO-Im three times a week^4^. During the indirect calorimetry study, mice were dosed with CDDO-Im on day 0 and 2. This resulted in an acute and significant decrease in food intake only at day 0. Unlike Shin et. al. where food intake was not measured during the 82-day experimental feeding, our study measured food intake once a week for 6 weeks. Hence, our findings demonstrate a long-term food intake reduction and prevention in weight gain in mice fed a HFD feed incorporated with CDDO-EA. Furthermore, it was not determined whether CDDO-Im prevented elevated blood glucose and plasma insulin levels in animals fed a HFD for 82 days^4^. We show that CDDO-EA protects from HFD-induced elevated blood glucose and plasma insulin levels. Our study also shows that these effects were not seen in mice fed a LFD into which CDDO-EA had been incorporated, demonstrating that CDDO-EA effects are specific to HFD feeding. Contradictory findings have been reported for CDDO-Me on its anti-obesity and anti-diabetic properties. Camer, et. al. fed C57BL/6J male mice a HFD (40% calories from fat) for 12 – 16 weeks and then orally gavaged them with CDDO-Me for two weeks. Although these mice showed lower fasting blood glucose and plasma insulin levels, they had similar food intake and body weight as the control obese mice fed a HFD and treated with vehicle ^5^. In other published reports, C57Bl6/J male mice were fed a HFD (40% calories from fat) and given an oral dose of CDDO-Me in the drinking water for 21 weeks^6, 7^. In this reported study, CDDO-Me reduced body weight in mice fed a HFD.

Although not consistent throughout the 21-week study, a reduction of energy intake was also observed during different weeks of the study in mice given CDDO-Me. Although these mice showed reduced body weight gain and food intake, CDDO-Me normalized glucose levels at 120 min in GTT following intraperitoneal injection of glucose. During the OGTT we performed, mice were orally gavaged with glucose, and we show that CDDO-EA normalized glucose levels starting at 60 minutes. Further, CDDO-Me lowered insulin levels in mice fed a HFD compared to HFD alone. This result is similar to our study in that CDDO-EA prevented an increase in fasting insulin levels in mice fed a HFD. The discrepancies in the animal studies could be due to the differences in the CDDO derivatives studied, method and duration of CDDO derivative administration, dose of CDDO derivative, percent of fat in diets, and sex of animals. Our present study demonstrates that incorporation of CDDO-EA in a HFD prevents development of insulin resistance and improves glucose homeostasis by inhibiting energy intake and obesity.

## AUTHOR’S CONTRIBUTIONS

AEC, SM, and SMR designed experiments, analyzed, and interpreted data, and wrote original draft; AEC, CR, SM, DC, IR, AAG, LRL, and SMR conducted experiments, collated, and interpreted data; JCLA interpreted data; AEC, CR, SM, JLCA, LRL, and SMR participated in the review and editing of the manuscript.

## CONFLICT OF INTERESTS

The authors declared no potential conflicts of interest with respect to the research, authorship, and/or publication of this article.

## ETHICAL APPROVAL

The animal study was approved by the Institutional Animal Care and Use Committee of The University of Texas Rio Grande Valley (Protocol # 17-05).

## FUNDING

The authors disclosed receipt of the following financial support for the research, authorship, and/or publication of this article: This work was supported in part by the National Institutes of Health [GM127272] and The University of Texas Rio Grande Valley Faculty Research Seed Grant and Institutional Seed Research Program Award.

## Notes

### Competing Interest Statement

The authors have declared no competing interest.

