## Supplemental Figure S1 - S3 for "Triterpenoid CDDO-EA Protects from Hyperglycemia, Hyperinsulinemia, and Obesity by Decreasing Energy Intake"

Supplemental Material

**Synthesis of CDDO-EA.**

Synthesized CDDO-EA was purified by column chromatography. The structure was confirmed with ^1^H NMR and HRMS, and the purity was verified by HPLC.

^1^H NMR spectrum was recorded on a Bruker 600 MHz spectrometer and chemical shifts are referenced to internal solvent resonances; multiplicities are indicated by s (singlet), d (doublet), t (triplet), q (quartet), m (multiplet) and br (broad). Coupling constants, *J*, are reported in Hertz. HPLC was performed using a ThermoFisher Vanquish with an Acclaim 120 C18 (5 μM, 4.6 x 100mm) column. Mass spectrum is obtained by Thermo Scientific QE Focus with atmospheric pressure chemical ionization (APCI).

^1^H NMR (600 MHz, CDCl3) δ 8.06 (s, 1H), 5.99 (s, 1H), 5.80 (t, 1H, *J* = 5.4), 3.27-3.40 (m, 2H), 3.08 (d, 1H, *J* = 4.5 Hz), 2.87 (d, 1H, *J* = 12.8 Hz), 1.99 (td, 1H, *J* = 13.8, 3.8 Hz), 1.77-1.85 (m, 5H), 1.68-1.76 (m, 1H) 1.52-1.68 (m, 6H), 1.50 (s, 3H), 1.35 (s, 3H), 1.28 (s, 3H), 1.30-1.39 (m, 2H) 1.19 (s, 3H), 1.15 (t, 3H, *J* =7.2 Hz), 1.04 (s, 3H), 1.02 (s, 3H), 0.93 (s, 3H);

HRMS (calcd. for C33H47N2O3 [M+H]^+^) 519.3581, found 519.3574

**Figure S1**. ^1^H NMR of CDDO-EA

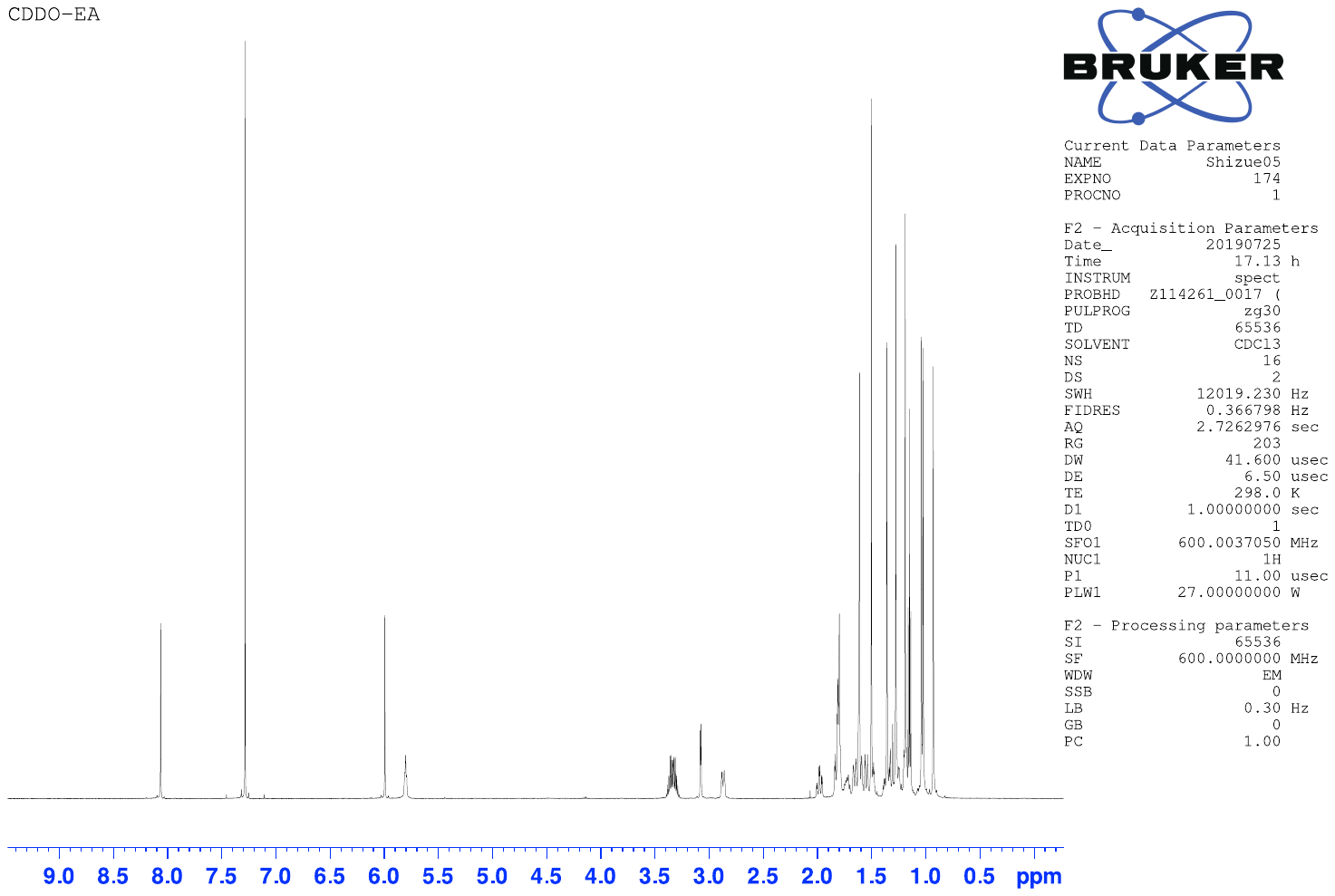

**Figure S2**. Mass spectrum of CDDO-EA (top) and calculated data (bottom)

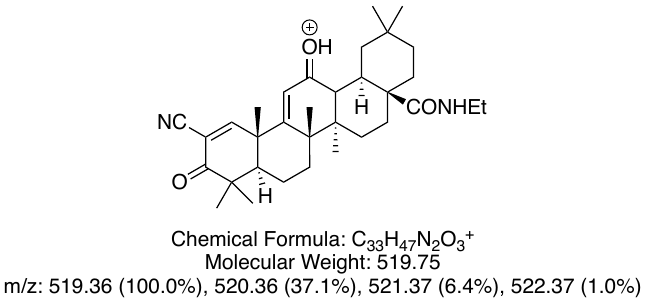

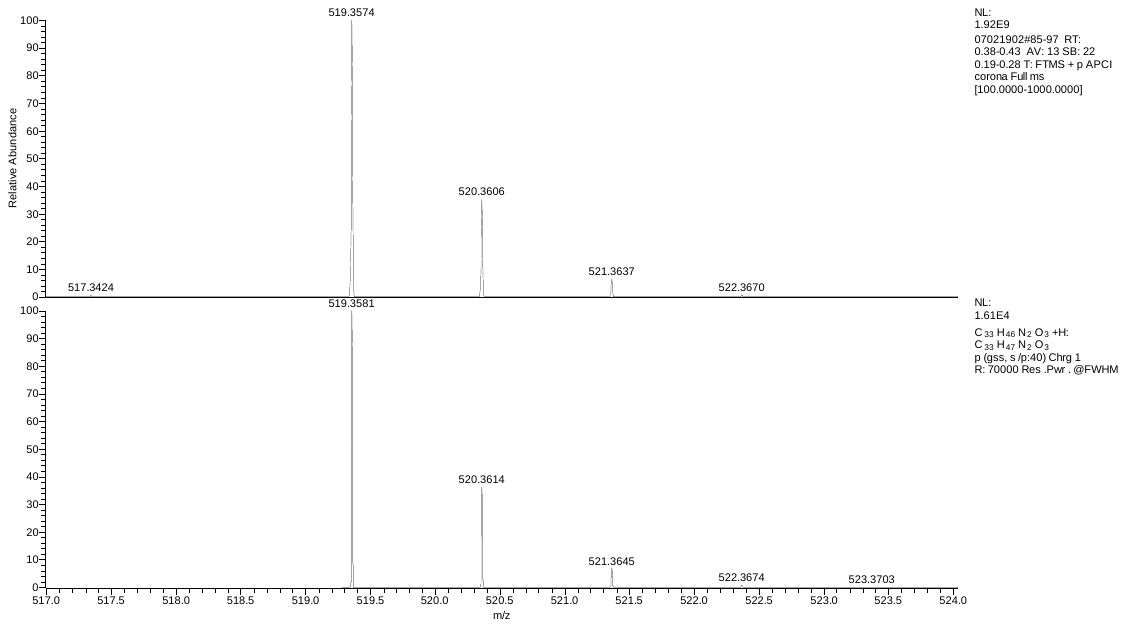

**Figure S3**. HPLC chromatogram of purified CDDO-EA: Flow rate 1.0 mL/min with MeOH.

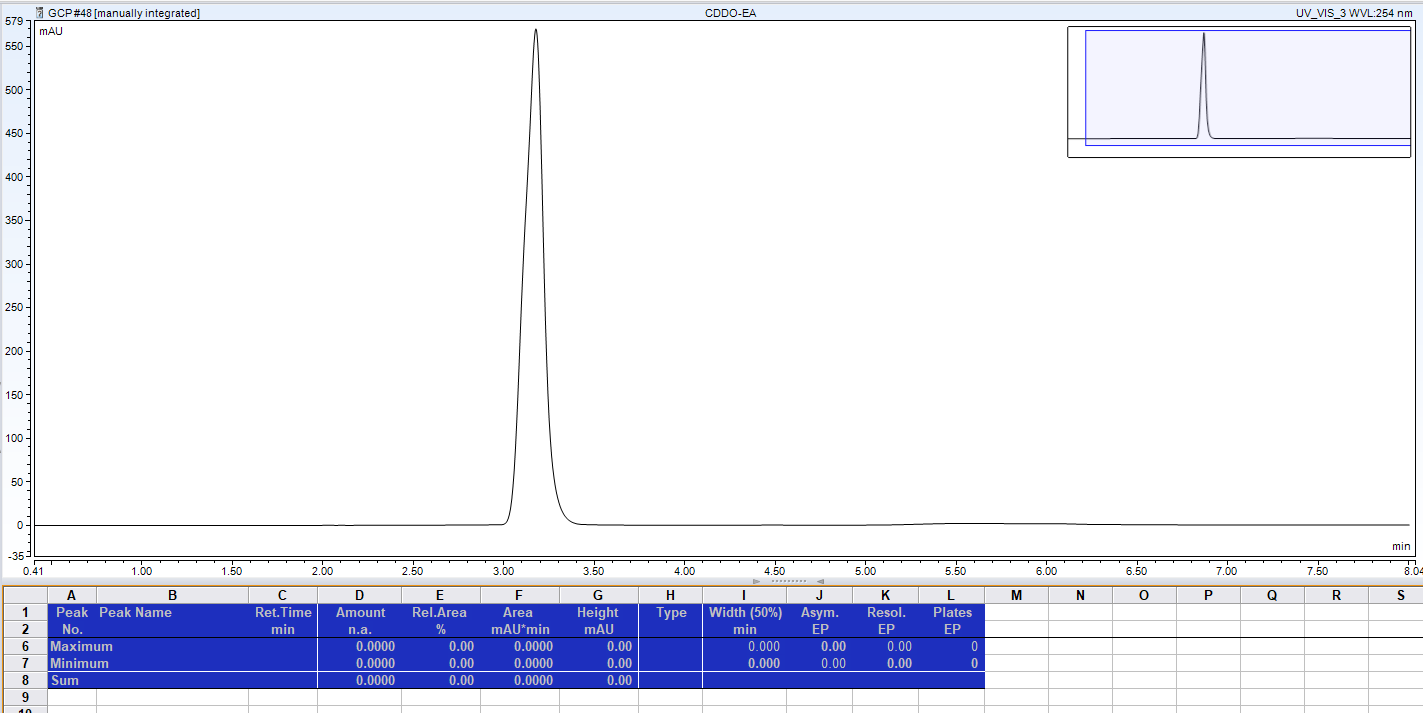
